# Ebola Virus VP35 Interacts Non-Covalently with Ubiquitin Chains to Promote Viral Replication Creating New Therapeutic Opportunities

**DOI:** 10.1101/2023.07.14.549057

**Authors:** Carlos A. Rodríguez-Salazar, Sarah van Tol, Olivier Mailhot, Gabriel Galdino, Natalia Teruel, Lihong Zhang, Abbey N. Warren, María González-Orozco, Alexander N. Freiberg, Rafael J. Najmanovich, María I. Giraldo, Ricardo Rajsbaum

## Abstract

Ebolavirus (EBOV) belongs to a family of highly pathogenic viruses that cause severe hemorrhagic fever in humans. EBOV replication requires the activity of the viral polymerase complex, which includes the co-factor and Interferon antagonist VP35. We previously showed that the covalent ubiquitination of VP35 promotes virus replication by regulating interactions with the polymerase complex. In addition, VP35 can also interact non-covalently with ubiquitin (Ub); however, the function of this interaction is unknown. Here, we report that VP35 interacts with free (unanchored) K63-linked polyUb chains. Ectopic expression of Isopeptidase T (USP5), which is known to degrade unanchored polyUb chains, reduced VP35 association with Ub and correlated with diminished polymerase activity in a minigenome assay. Using computational methods, we modeled the VP35-Ub non-covalent interacting complex, identified the VP35-Ub interacting surface and tested mutations to validate the interface. Docking simulations identified chemical compounds that can block VP35-Ub interactions leading to reduced viral polymerase activity that correlated with reduced replication of infectious EBOV. In conclusion, we identified a novel role of unanchored polyUb in regulating Ebola virus polymerase function and discovered compounds that have promising anti-Ebola virus activity.

**Significance Statement:** Ebola virus infection can result in high mortality rates with extreme risk of person-to-person transmission. Sporadic outbreaks in Africa have resulted in thousands of fatal cases, highlighting that there is still insufficient knowledge to develop effective antiviral therapies. Like other viruses, Ebola utilizes the host machinery to replicate. Understanding how viral and host proteins interact can help identifying targets for the rational design of antiviral drugs. Here, we identified interactions between the cellular ubiquitin machinery and the Ebola virus polymerase cofactor protein VP35, which are important for efficient virus replication. We developed an approach to identify and block these virus-host interactions using small chemical compounds, which provides a useful tool to study functional molecular mechanisms and at the same time a potential approach to antiviral therapies.

## Introduction

Ebola virus disease (EVD), characterized by severe hemorrhagic fever, is caused by the highly transmissible and lethal *Ebolavirus* (EBOV). The most pathogenic member of this family is *Zaire ebolavirus* (ZEBOV), which has been responsible for the highest number and most devasting outbreaks in Africa. EBOV is a non-segmented, negative-sense RNA virus from the genus *Ebolavirus* (family *Filoviridae)*. The viral RNA genome is encapsidated by the nucleoprotein (NP), which binds to the RNA-dependent RNA polymerase (RDRP) complex and transcription activator (VP30). The RDRP is composed of the catalytic subunit of the polymerase L, and the polymerase cofactor protein VP35, which interact with the viral envelope through the VP40 and minor VP24 matrix proteins to form the virion (1–3). In addition to VP35’s critical role as a cofactor of the viral polymerase, it has been extensively studied for its function in the inhibition of innate signaling pathways and antagonism of antiviral type-I Interferon (IFN-I) (4–15).

EBOV infection is initiated upon the virus binding to cell receptors via the viral glycoprotein (GP). EBOV enters the host cells either by pinocytosis, Clathrin-mediated, or caveolin-mediated endocytosis (1, 16, 17). Once the viral genome is released into the cytoplasm, transcription, and replication occur through a positive sense RNA (antigenome) and is carried out by the viral RDRP. VP35 can interact with several host proteins as well the viral polymerase to promote virus replication and do so either via viral polymerase activity or by antagonizing immune signaling (4, 6, 12, 14, 15, 17, 18). VP35 also is known to inhibit IFN-I production by targeting the pattern recognition receptor RIG-I (12). The C-terminal region of VP35, termed the IFN-inhibitory domain (IID), inhibits IFN induction by sequestering the viral 5’-triphosphorylated dsRNA from RIG-I (8, 12, 19, 20). Extensive previous structural and functional work on the IID of VP35 identified two basic regions; a basic patch comprised of residues K222, R225, K248, and K251, involved in viral polymerase activity, and a central basic patch (R305, K309, R312, R319, R322, K339) with a major function in IFN inhibition (5, 8, 12, 19, 20), although it can also affect polymerase activity (6, 11). Importantly, mutations disrupting the basic charge on R225 decreased VP35 polymerase cofactor activity, and this could be in part due to loss of interaction with NP (11). However, the precise mechanism remains unclear. The different functions of VP35 have been the focus of much research for the development of treatments and vaccines.

The process of viral replication and particle release can be regulated by post-translational modifications on different viral proteins, which additionally can alter several intracellular antiviral mechanisms (4, 21–30). The process of ubiquitination has been found to enhance virus replication by direct ubiquitination of viral proteins or indirectly by counteracting host defenses or modulating the innate immune response (4, 21–26). For example, in terms of direct modulation of virus replication, the HECT E3 ubiquitin (Ub) ligases SMURF2, Nedd4, ITCH, and WWP1 can promote ubiquitination of VP40 for efficient virus budding and virus-like particle (VLP) egress (24, 26–31). In addition, VP40 stability can be regulated by modification with SUMO (for Small Ub-like Modifier) (22), which can have a similar function to ubiquitination. We also recently reported that ubiquitination on VP35 is required for optimal viral transcription by promoting efficient interactions with the viral polymerase L and thus plays a critical role in viral pathogenesis (4). VP35 is covalently ubiquitinated on lysine 309 (K309) by the host E3 ubiquitin ligase TRIM6, which is crucial for efficient viral replication (4). TRIM6 depletion or a recombinant EBOV VP35 - K309R mutant virus lacking ubiquitination on K309, results in reduced viral transcription and disrupted production of viral proteins. Moreover, a mutant of VP35 lacking ubiquitination on K309 interacts with higher affinity with the viral nucleoprotein resulting in dysregulated virus assembly (6). We also previously found that VP35 can bind Ub via non-covalent interactions; however, the precise function of these VP35-Ub interactions remains unknown (4, 6). Unanchored polyubiquitin (polyUb) chains have been shown to play a role in immune signaling (32–36), and may also play a role in virus uncoating (37). However, in our knowledge, no role has yet been identified for unanchored Ub in viral polymerase function. Here, we show that VP35 interacts with unanchored K63-linked polyUb chains, promoting efficient viral polymerase function and EBOV replication. A VP35 R225E mutant, which was previously reported to have reduced binding with NP and reduced activity in minigenome assays (11), showed reduced binding to unanchored Ub, providing a potential explanation for the reduced polymerase activity. We identified novel antiviral compounds that block non-covalent interactions between VP35 and Ub, providing a potential novel approach to the development of antiviral strategies.

## Results

### The C-terminal IID of VP35 Interacts with Unanchored K63-linked polyubiquitin Chains

We previously reported that VP35 is covalently ubiquitinated on K309 using co-immunoprecipitation (co-IP) assays. Intriguingly, these experiments also consistently showed a non-modified fraction of VP35 that co-immunoprecipitated with Ub, suggesting a non-covalent interaction between Ub and VP35 (4). In this new work, we postulate that VP35 interacts with unanchored or free polyUb chains and that the interaction between these Ub chains and VP35 is functionally relevant. Here, to first confirm binding between VP35 and Ub, we performed a co-IP assay in which we pulled down ectopically expressed wildtype (WT) Ub or a Ub mutant lacking the C-terminal di-glycine residues (HA-Ub-ι1GG), which renders Ub unable to form covalent linkages. The terminal -GG on Ub is required for the formation of covalent conjugates of Ub with other proteins (38, 39). This approach allows testing non-covalent interactions between monomeric, non-conjugated Ub, and VP35. Consistent with our previous observation, in the presence of WT Ub, multiple migrating forms corresponding to the molecular weight of covalently ubiquitinated VP35, as well as monomeric VP35 (non-covalent interaction with Ub), were detected by immunoblot (IB). In contrast, monomeric, non-conjugated VP35, co-immunoprecipitated with HA-Ub-ι1GG (Figure 1A, IP). As expected, the HA-Ub-ι1GG runs at the predicted molecular weight of monomeric Ub (∼8.5 kDa) and is unable to form the characteristic smear corresponding to cellular ubiquitinated proteins (Figure 1A, Whole Cell Extract [WCE]). This indicates that VP35 associates non-covalently with Ub, either directly or indirectly.

**Figure 1.**
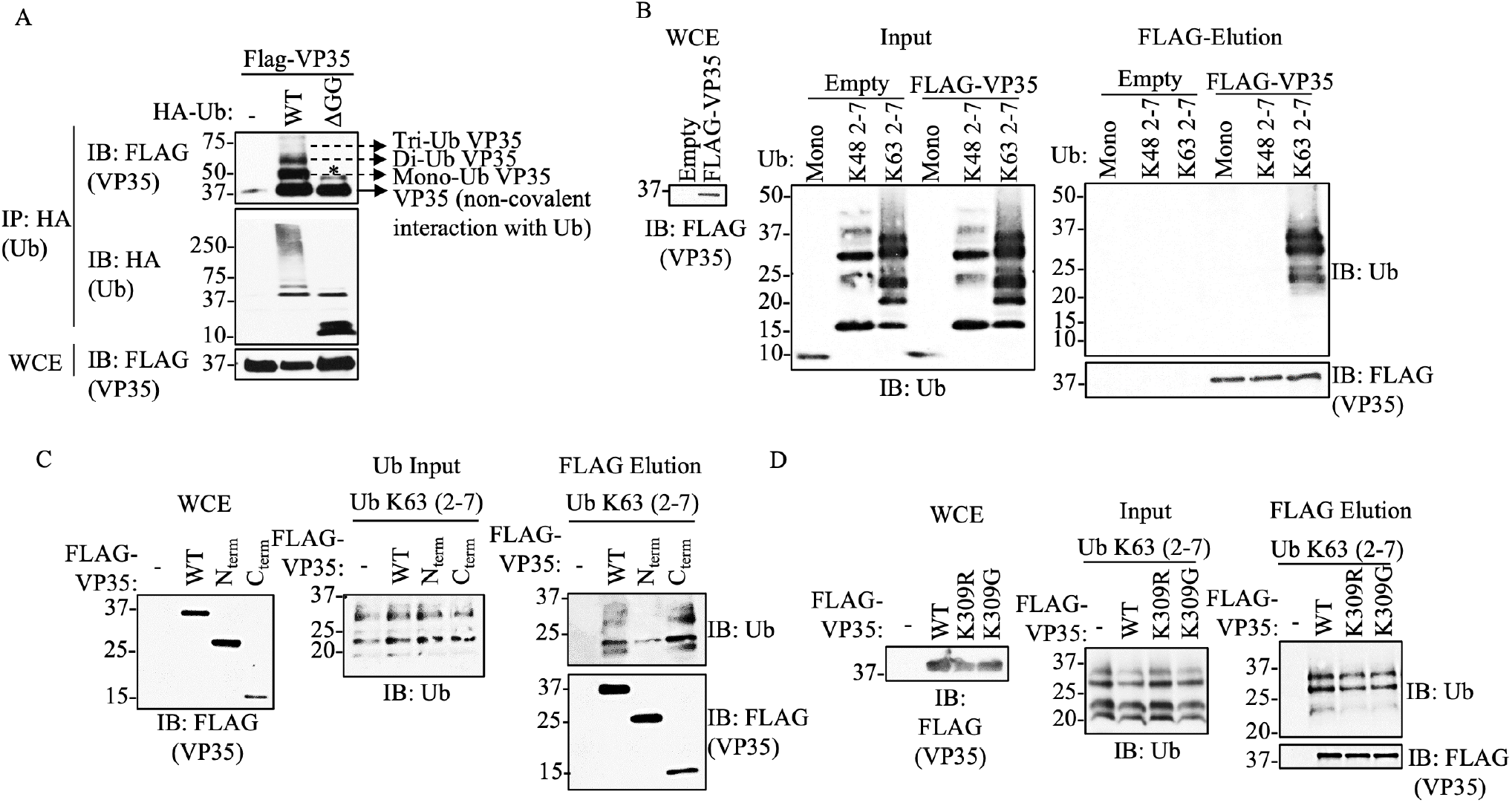
EBOV VP35 protein interacts with Ub non-covalently. A) Whole Cell extracts (WCE) from HEK293T cells transfected with Flag-VP35 (VP35) and HA-Ub wild type (WT) or HA-Ub ΔGG (cannot conjugate proteins), were used for HA immunoprecipitation (IP) under non-denaturing conditions (RIPA washes), followed by immunoblot (IB). B) Purified recombinant K48 or K63 polyUb chains (mix of 2-7 Ub chains), were mixed in vitro with Flag-VP35, followed by Flag IP. Interacting proteins were eluted with Flag peptide. C-D) Experiments performed as in (B) but using the C-terminal IID domain of VP35 (C), or the VP35 K309R or K309G mutants, which are not covalently ubiquitinated (D). * No ubiquitinated VP35, possibly phosphorylation.

To further confirm direct, non-covalent, binding between Ub and VP35 and to identify the type of polyUb chains involved in these interactions, a cell-free *in vitro* binding assay with purified Flag-VP35 and recombinant purified unanchored K48- or K63-Ub chains (a mix of 2-7 Ub chains) was conducted. We found that VP35 strongly interacts with unanchored K63-but not K48-Ub chains (Figure 1B). Furthermore, unanchored K63-polyUb chains interacted mostly with the C-terminal IID of VP35 (Figure 1C). Finally, since we previously showed that VP35 K309R or K309G mutants lose covalent ubiquitination on the K309 residue (6), we asked whether these mutants can still bind unanchored Ub. We found that these mutants interacted with unanchored Ub at similar levels compared to VP35 WT (Figure 1D), suggesting that non-covalent interactions with Ub do not require covalent ubiquitination on the K309 residue.

### Unanchored Polyubiquitin chains promote Ebola virus polymerase activity

Since we previously found that ubiquitination on the VP35 IID domain can regulate polymerase activity, we asked whether unanchored Ub would also affect this function. To test the function of unanchored Ub in relation to VP35 polymerase co-factor activity, we used the unanchored Ub-specific protease Isopeptidase T (IsoT, also called USP5), which can cleave unanchored Ub by interacting with the free di-glycine residue of Ub chains (40). Ectopic expression of IsoT-WT cleaved polyUb chains, as observed in WCE, and correlated with reduced association of Ub with VP35 (Figure 2A). In contrast, a catalytically inactive mutant (C335A), which does not cleave unanchored Ub (40), did not affect the association between VP35 and Ub chains (Figure 2A). These results support that VP35 interacts with unanchored Ub chains. Furthermore, the effects of IsoT correlated with decreased EBOV polymerase activity evaluated in a minigenome assay (Figure 2B), suggesting that unanchored Ub may promote virus replication.

**Figure 2.**
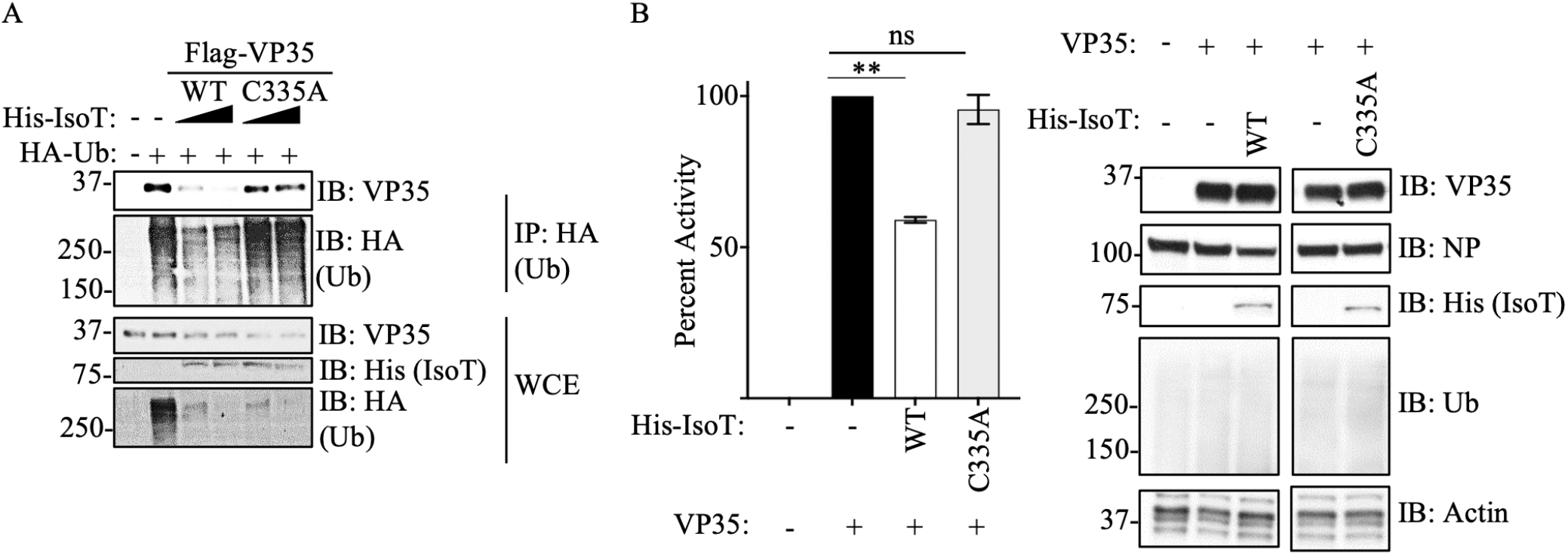
Unanchored Ubiquitin interactions with VP35 promote viral polymerase activity. **A)** WCE from HEK293T cells transfected with His-IsoT WT, His-IsoT C335A, VP35 WT, and HA-Ub were used for IP with anti-HA beads. B) Polymerase minigenome assay. HEK293T cells transfected with a monocistronic firefly luciferase-expressing minigenome, including VP30, L, and REN-Luc/pRL-TK, in the presence or absence of IsoT-WT or C335A mutant. Data are expressed as Mean + SEM of three independent assays in triplicate. Tukey’s multiple comparisons tests. ** p < 0.001. The percent of activity from the luciferase and renilla (Luc/ren) ratio was calculated.

### Identification of amino acid residues involved in Ubiquitin – VP35 interactions

We employed a computational approach to identify specific amino acids in contact between VP35 and Ub. We first predicted the structure of the C-terminal IID domain of VP35 in complex with Ub, using a combination of protein docking and molecular dynamics simulations (Figure 3A).

**Figure 3.**
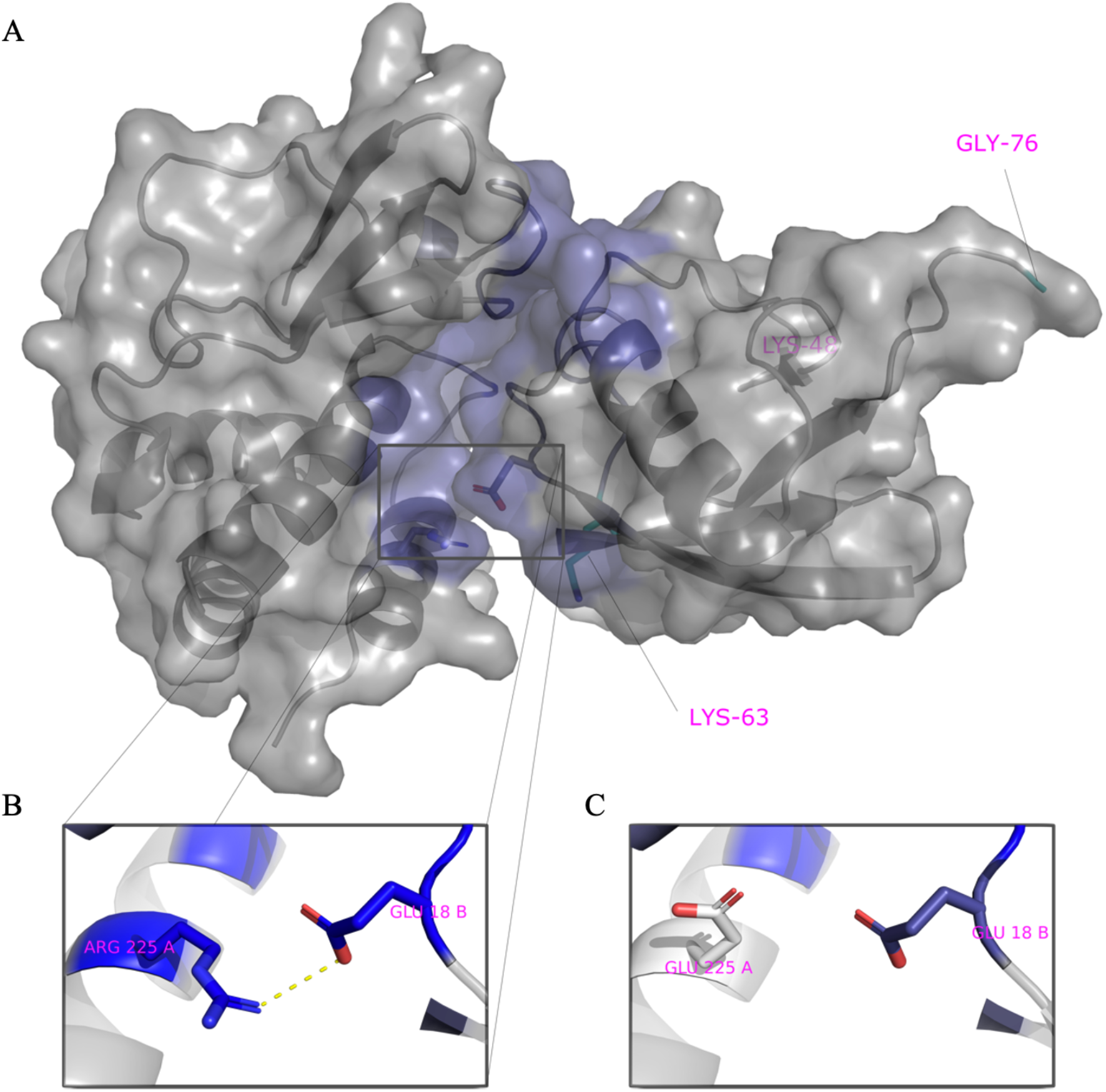
Model of VP35 interacting with Ubiquitin. A) The complex of VP35 (PDB ID 3JKE) and Ubiquitin (PDB ID 1UBQ) modelled using a combination of protein docking and molecular dynamics simulations. Within the complex, VP35 is shown on the left and Ubiquitin on the right. The K48 and K63 Ub residues are shown in cyan on the bottom left and C-terminal on the right within Ub. B) One of the strongest interactions contributing to the stability of the complex is ARG225-GLU18. C) Mutation of ARG225 to GLU affects interactions.

We utilized the Surfaces software to determine the top contributing interactions between VP35 and Ubiquitin. This analysis detected the R298-E24, R225-E18, R305-S57, R305-D58 and Y229-E18 as the top contributions to the VP35-Ub interaction (Figure 3A and Supplementary Table 1). We utilized gRINN to validate the Surfaces result. The gRINN analysis confirmed all five interactions as the top contributions (See Supplementary Table 1). Both methods suggest that R225-E18 is among the top contributors to the interaction. Analysis of all possible mutations at position R225 with Surfaces (Supplementary Table 2), suggested that the R225E mutation in VP35 would abrogate the interaction (Figure 3B-C). The R225E mutation was experimentally tested and has functional effects, abrogating polymerase activity in minigenome assays (Figure 4A), which is consistent with previous studies (11). Importantly, the mutation led to a decrease in K63 polyUb binding in a cell-free in vitro coIP assay (Figure 4B), or by mixing lysates from cells expressing VP35 and Ub (Figure 4C). In contrast, the mutation R225K, which maintains the positive charge on this residue and was predicted to not completely disrupt the interaction (Supplementary Table 1), partially rescued binding with Ub (Figure 4C). Therefore, the reduced binding of VP35-R225E and its reduced polymerase activity further supports a functional role for non-covalent interactions between VP35 and Ub in promoting viral polymerase activity and suggests that the modelled complex structure is correct.

**Figure 4.**
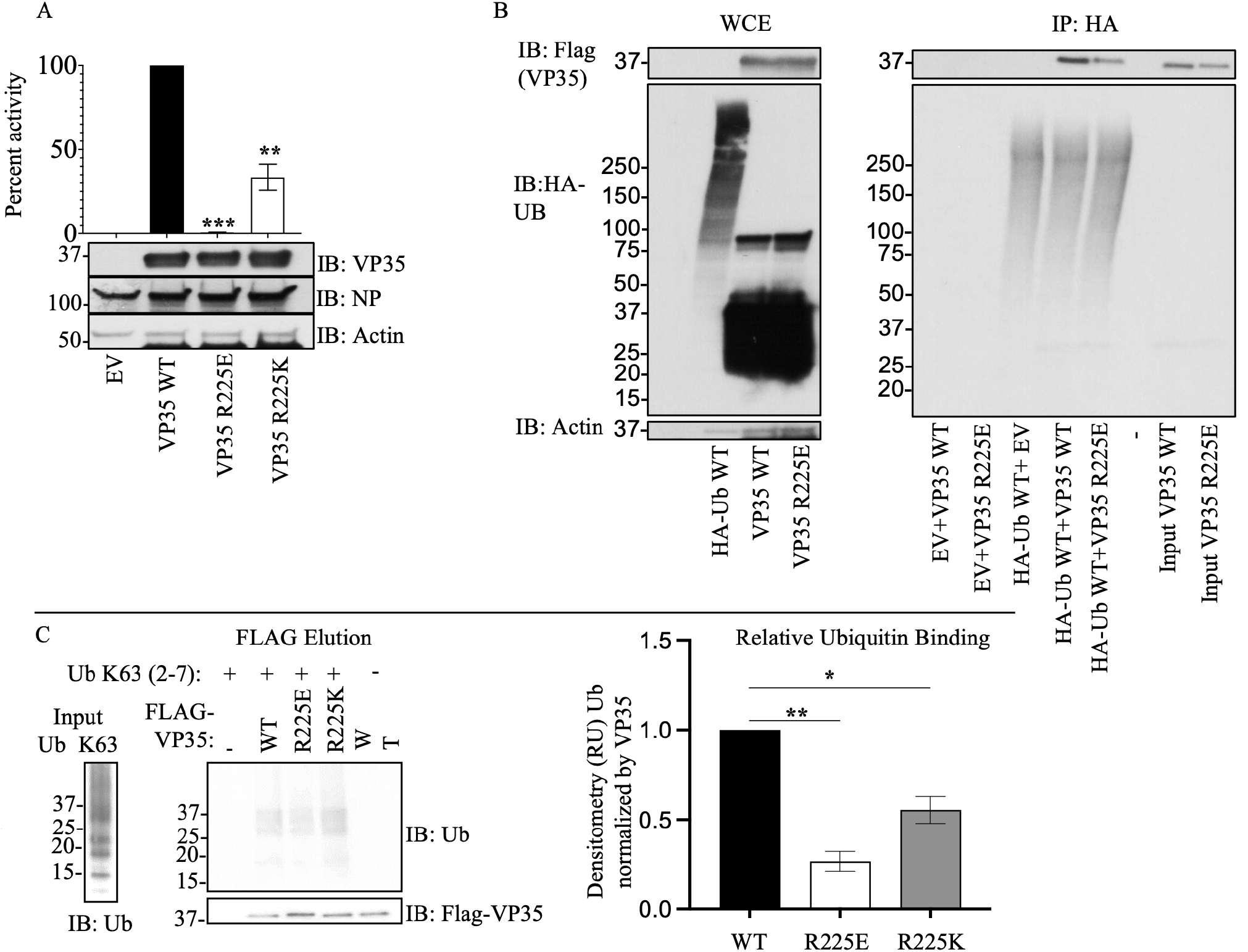
An intact R225 residue on VP35 is required for optimal interaction with Ub and viral polymerase function. A) HEK293T cells were transfected with minigenome plasmids and VP35 WT, VP35 R225E, or VP35 R225K, followed by Luciferase assay. B) HEK293T cells were transfected with plasmids encoding Flag-VP35 WT, VP35 R225E, or VP35 R225K. WCE were then used to isolate Flag-tagged proteins using anti-Flag beads. After washes, the beads containing VP35 were mixed with the WCE containing HA-Ub to test binding. C) as in B, but instead of mixing with WCE, binding was performed using purified recombinant K63-linked polyUb chains, followed by Flag elution. Quantification by densitometry of 3 independent experiments is shown.

Since our model indicates that Ub interacts with the basic patch of VP35 that modulates polymerase activity, it would not be likely that Ub affect dsRNA binding to VP35, based on previous reports (5, 11). To test this possibility, we used the core model in Figure 3A as a template to create a model of the ternary complex of VP35, dsRNA and a tri-Ub chain of K63-linked polyubiquitin (Figure 5 and Supplementary PyMol session file 1). Interestingly, in this model K48 PolyUb would clash with the dsRNA binding site while the position of K63 points away from the dsRNA binding site (Figure 5A), in agreement with the experimental data (shown in Figure 1). The model complex suggests that the central Ub bound to VP35 makes contact with RNA. Not only is K48 occluded by the RNA in the model but it in fact makes favorable interactions with the RNA (Figure 5B). Using Surfaces to identify per-residue contributions to this extended interface with RNA, we identified that K48, R54, Y59, and A46 among others favorably contribute to binding RNA. These contribute to strengthen the overall estimated binding free energy by 25% relative to that of the interface between VP35 residues and RNA but lacking the interaction with Ub. Therefore, the interaction between Ub and RNA is likely to be functionally important and suggests that VP35-Ub interaction does not affect the ability of VP35 to bind dsRNA and antagonize IFN-I.

**Figure 5.**
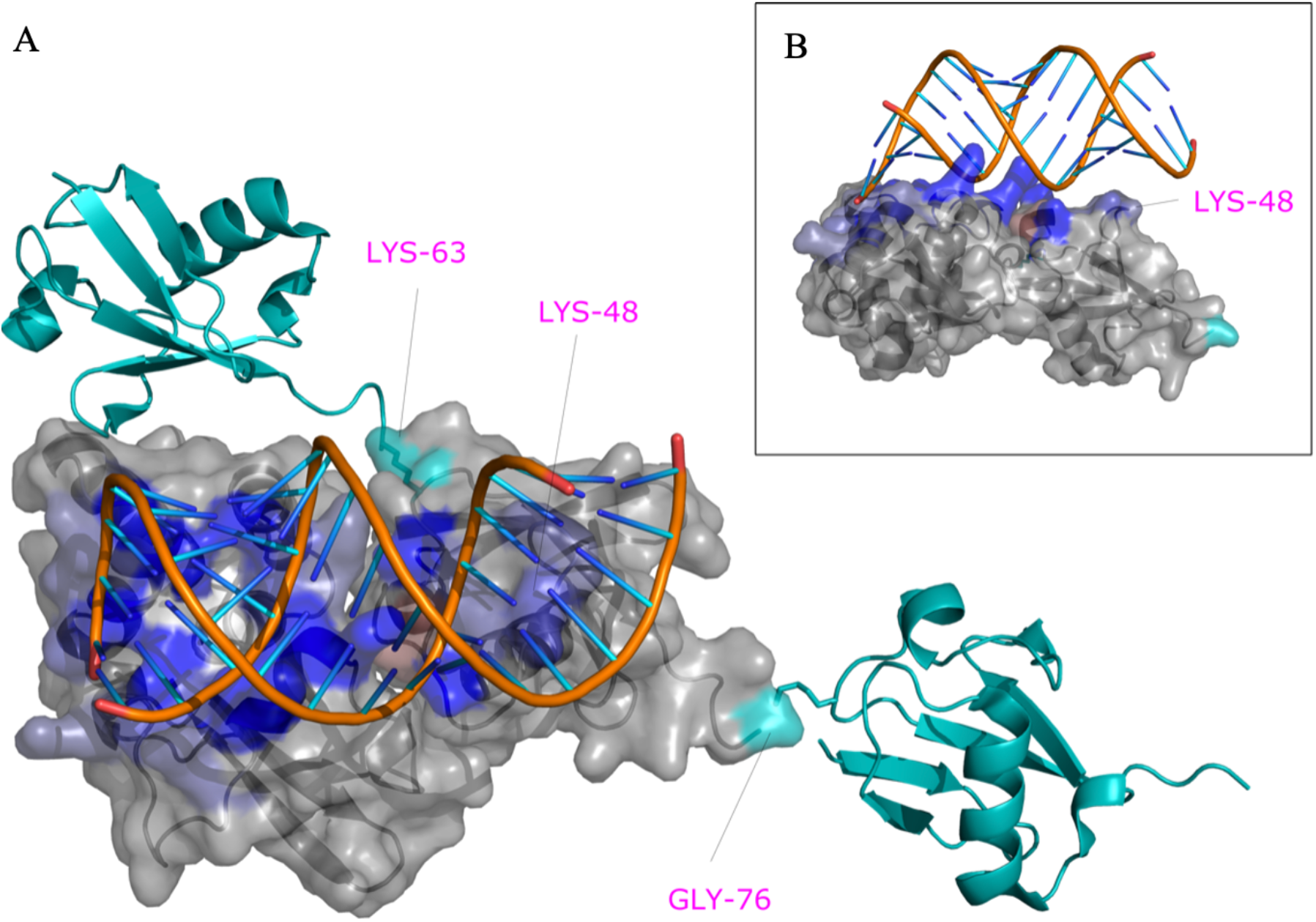
VP35-PolyUb-dsRNA predicted complex. The predicted structure of the VP35-Ub complex was used as a template to superpose the structure of VP35 bound to RNA (PDB ID 3KS8). PolyUb was modeled using as a template the structure of K63 Di-Ubiquitin (PDB ID 2JF5). The residues K63, K48, and G76 of the central Ub bound to VP35 are labeled in magenta and contribute favorably to RNA binding in this model.

### Identification of small-molecules that inhibit VP35 – unanchored K63 Ub Interactions

To test whether the non-covalent interactions between VP35 and Ub have functional relevance, we first employed a computational approach with the objective of identifying compounds that could potentially disrupt the Ub-VP35 complex. A cavity within the putative VP35-Ub interface was used as a target to dock 36,000 small molecules with known complex structures using the small-molecule protein docking program FlexAID. Two criteria were used to detect potential binders: a combination of highly favorable docking score relative to the average of all molecules and a large level of binding-site similarities measured using the IsoMIF program between the targeted VP35 cavity and the original protein where the compound is known to bind (Figure 6A). The docking scores (CF) for the 36,000 molecules had a mean value around -100 AU. The z-score of the top 10% varied from -5.0 to -8.0. The top-scored molecules were evaluated to identify those molecules among the top 10% likely to have favorable pharmacological properties. Two molecules emerged from this analysis, pCEBS, 3-[4-(aminosulfonyl) phenyl] propanoic acid – a molecule developed to inhibit carbonic anhydrase (41), and SFC, 2,5-dimethyl-4-sulfamoyl-furan-3-carboxylic acid – a molecule developed as a Metallo-Δ-lactamase inhibitor (42). The two candidates pCEBS and SFC, had a CF value of -321AU and -278AU, equivalent to a Z-score of -6.5 and -4.8, respectively. The binding site analysis with IsoMIF of the cavities of the crystal structures of the complexes containing SFC (PDB: 6KXO, 6KXI, and 6LBL) revealed binding site similarities of 0.25, 0.32, and 0.35 with VP35, respectively, and the cavities of the crystal structures of the complexes containing pCEBS (PDB: 2NN0 and 2NN1) showed binding site similarities of 0.24 and 0.28 respectively to VP35. The mean binding site similarity for the top 10% of molecules in the docked dataset is 0.21. Thus, the chosen two molecules have a docking score considerably lower (more favourable) than the average and the binding sites known to bind these molecules are more similar to the targeted VP35 cavity than cavities of other top-scoring molecules. This suggests that important interactions responsible for binding pCEBS and SFC are also exploited in the VP35 cavity. Although independently selected, the two compounds share a common sulfonamide group (-S0_2_NH_2_) linked to an aromatic ring system and a carboxyl group that interacts with the same VP35 residues and both are nearly perfectly superimposed (Figure 6B-C). Interestingly, at least one X-ray structure of VP35 (PDB ID 4IBG) shows a sulfate ion from the crystallization buffer bound in very close proximity to the position where the sulfonamide group from pCEBS and SFC are predicted to interact with VP35 based on the docked ligand poses (Figure 6D). The experimental observation that a sulfate ion at that position has favourable interactions with VP35 serves as indication that the docked structures with their sulfonamide groups located at that approximate position are taking advantage of interactions that were experimentally validated.

**Figure 6.**
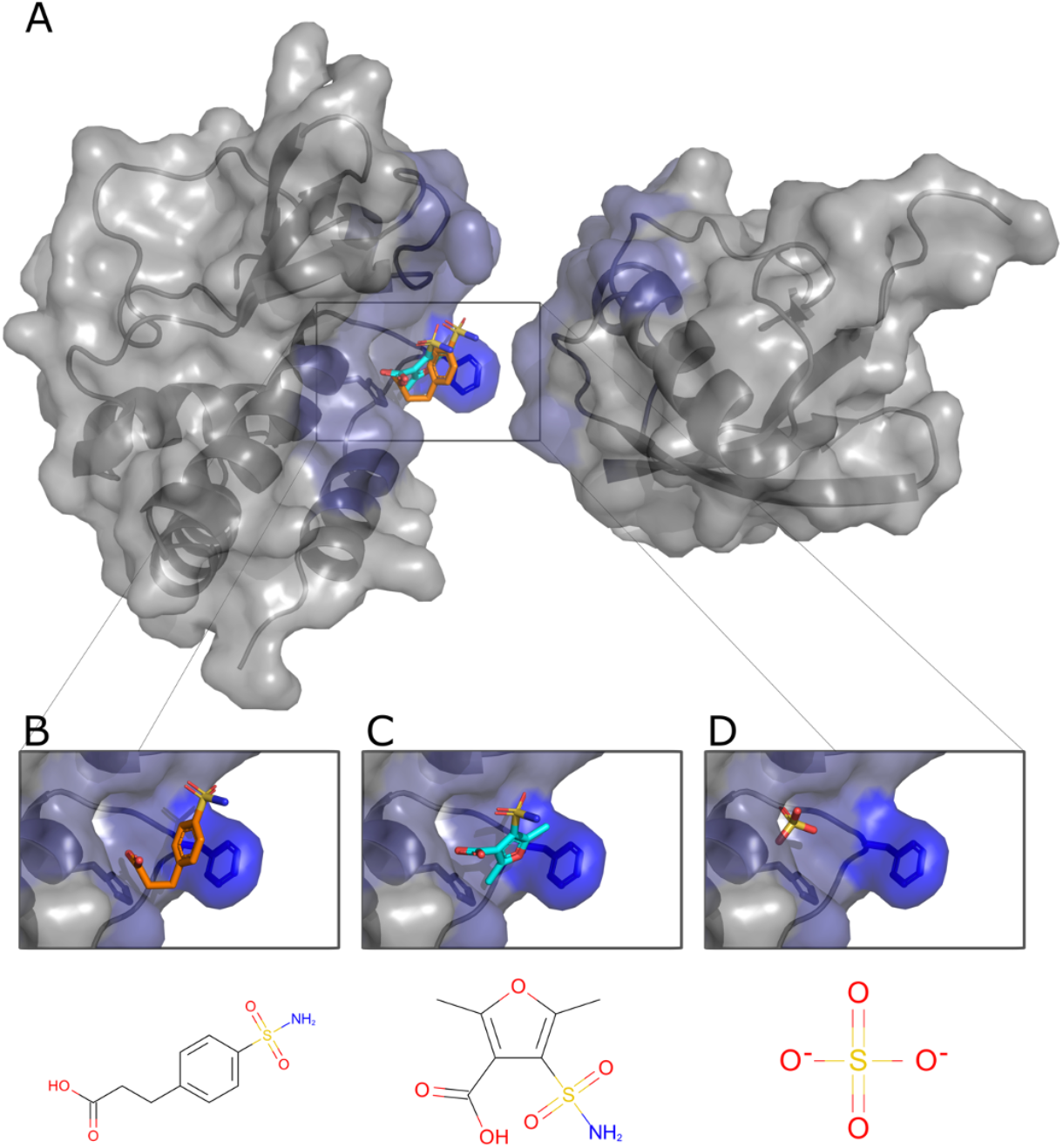
Predicted poses of small-molecules disrupting the VP35-Ub interaction. A. The cavity within the interface with Ub with predicted bound pCEBS (B) and SFC (C) in close proximity to a sulfate ion (D) observed experimentally (PDB ID 4IBG).

Incubation with the two highest concentrations of either of the two molecules leads to a decrease in Ub-VP35 interactions detected in coIP assays (Figure 7A), and these correlated with a decrease in luciferase activity in minigenome experiments (Figure 7B). Although the inhibition did not show a perfect dose-response, possibly due to other important interactions between Ub and viral polymerase proteins, these results, at high concentrations, further suggest that Ub-VP35 non-covalent interactions may contribute to efficient EBOV polymerase function. The compounds showed less than 5% cell death in a cytotoxicity assay (Figure 7C). Importantly, both molecules lead to a decrease of infectious ZEBOV replication in cells, as observed in plaque reduction (PR) and virus yield reduction (VYR) assays, (Figures 7D and 7E). The effect of pCEBS and SFC are within the same range as that observed for the nucleoside analog Favipiravir (T-705), used as a positive control for its broad-spectrum reported activity against Filoviruses (43). Taken together, these results suggest that VP35 interaction with Ub promotes EBOV replication by enhancing the function of VP35 as a co-factor of the polymerase. Furthermore, the identification of chemical compounds that block these VP35-Ub interactions could serve as starting point for the development of novel antivirals.

**Figure 7.**
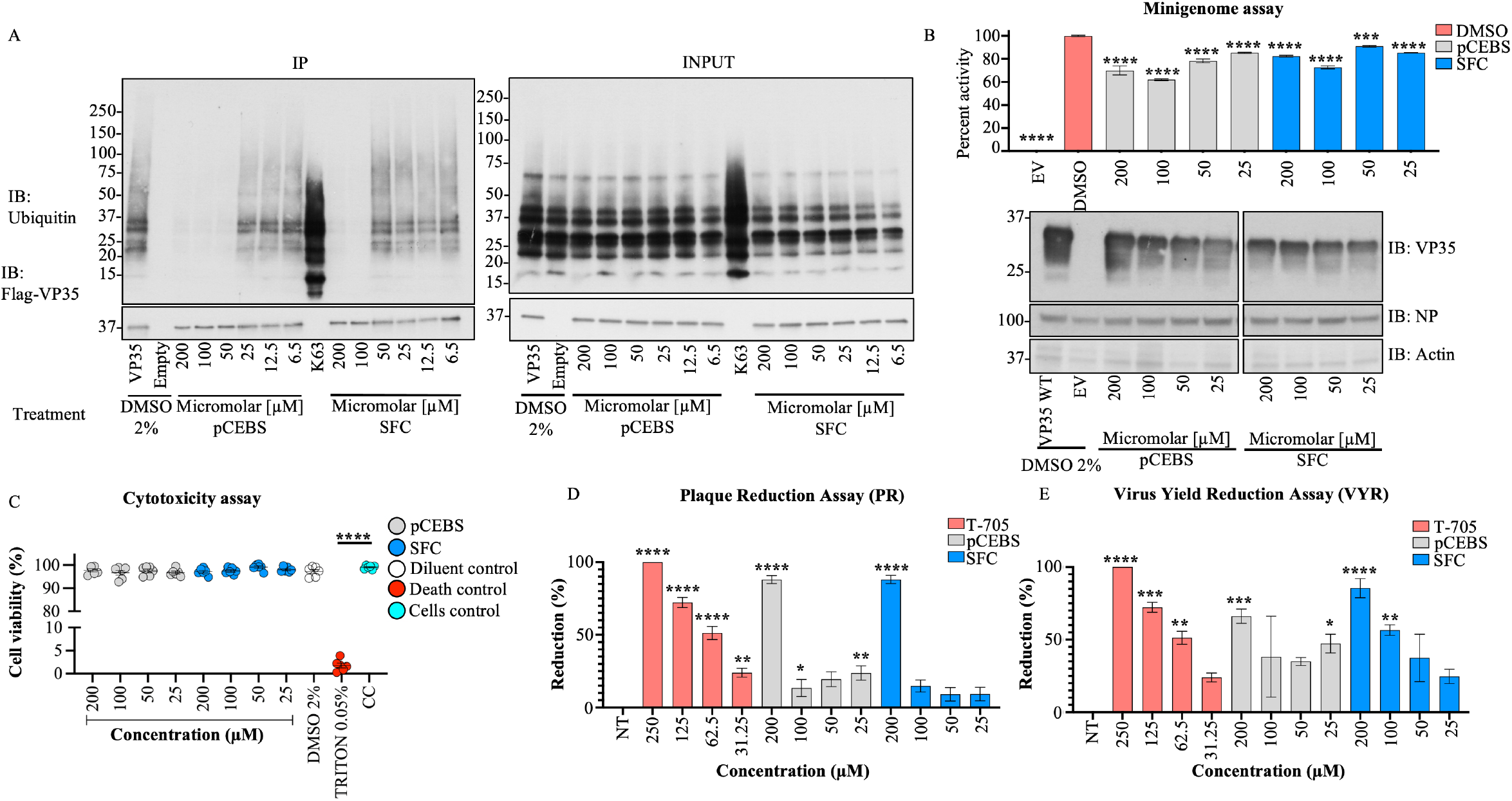
pCEBS and SFC compounds inhibit interactions between VP35 and K63-linked polyubiquitin chains and correlate with loss of viral polymerase activity and virus replication. A) Flag-VP35 bound to anti-Flag beads were incubated for 1h at room temperature with different concentrations of pCEBS or SFC, followed by incubation with recombinant purified unanchored K63-linked polyUb chains (2–7). VP35-Ub complexes were eluted with Flag-peptide and analyzed by Immunoblot. B) 293T cells were transfected with minigenome components and 4 hours post-transfection cells were treated with pCEBS and SFC compounds at different concentrations. 50 hours later cells were lysed for luciferase assay. C) Cytotoxicity test (CyQUANT MTT Cell Viability Assay ThermoFhisher) using pCEBS and SFC at different dilutions D) Plaque reduction and (E) Virus Yield Reduction assays, the cells were infected by 1 hour and after 1h the treatment was made with pcEBS, SFB compound or DMSO: Dimethyl sulfoxide with the overlay. The number of plaques in each set of compound dilution were converted to a percentage relative to the untreated virus control. The percent of activity from the ratio of luciferase and renilla (Luc/ren) was calculated. Data are depicted as Mean + SEM of the two independent assays in triplicate. Tukey’s multiple comparisons tests. p < 0.001 **, p < 0.0001 ***, p < 0.00001 ****.

## Discussion

In this study, we demonstrated that the VP35 protein of EBOV interacts non-covalently with unanchored K63-linked polyUb chains. This interaction between Ub and VP35 promotes viral polymerase activity leading to optimal virus replication. Our findings indicate that these Ub chains play a functional role in cells and *in vitro*, and this is supported by the following lines of evidence: (i) identification of specific residues on VP35 in contact with Ub, (ii) mutations that change the basic amino acid on VP35 R225 reduce interactions with Ub and correlate with reduced polymerase activity, (iii) mutation on K309, which is the acceptor of covalent ubiquitination, does not affect interaction with Ub, (iv) ectopic expression of IsoT, which specifically degrades unanchored Ub, reduced viral polymerase activity and VP35-Ub binding, and (v) chemical compounds predicted to block VP35 interactions with Ub reduce minigenome polymerase activity and infectious EBOV replication.

Using a pharmacological approach to block Ub-VP35 interactions we showed that these interactions are relevant in promoting polymerase activity and replication of infectious EBOV. However, loss of Ub interaction with VP35 did not show a perfect correlation with the reduced polymerase activity and virus replication. In addition, the compounds did not show a linear dose response demonstrating the challenges of inhibiting protein-protein interfaces (44), particularly in this case where it is possible that any of the Ub units within the Ub chain have additional potential interactions, not only with RNA as modelled here for the central Ub, but also with the N-terminal region of VP35 or other viral proteins. Furthermore, it is also possible that pCEBS and SFC have additional cellular targets that affect the minigenome assays. Alternatively, Ub interactions with VP35 could affect protein oligomerization or formation of large molecular complexes, and breaking these interactions with pCEBS and SFC could have larger effects on virus replication. For example, increasing amounts of VP35 result in a “bell curve” observed in minigenome assays (4, 45, 46), and could potentially be due to oligomerization and/or aggregation. Although we cannot rule out indirect effects of the compounds on other cellular or viral proteins at this stage, our data using VP35 mutants and small molecules that block Ub interactions with VP35 suggest that unanchored Ub plays a role, at least in part by promoting viral polymerase function.

The two compounds pCEBS and SFC add to the list of existing compounds that target VP35 activity. As the emergence of resistance mutations within viral proteins is common, the identification of novel compounds that likely interact with VP35 at a different binding-site than previously exploited (47), contributes to the future development of novel antivirals.

VP35 has been extensively studied mostly as a major IFN-I antagonist, and its role as a co-factor of the viral polymerase is less understood. VP35 also participates in nucleocapsid packaging and efficient viral replication. Previous studies have shown that the R225 amino acid is located on a second basic patch on VP35, consisting of residues K222, R225, K248 and K251 (5). Amino acids in this basic patch most likely do not contribute to VP35 binding to dsRNA (5), and therefore should not play a major role in IFN antagonism consistent with our computational model. In fact, the model in the presence of Ub and dsRNA suggests that the non-covalent interactions with Ub may even enhance IFN antagonism. However, the R225E mutation on VP35 reduces its activity in minigenome assays, most likely due to reduced interaction with NP, and not a loss of interaction with L (11). Our model suggests that VP35-R225 is in contact with Ub and reduced interaction with Ub may explain the reduced minigenome activity previously observed. Although we did not test whether the compounds have an effect on IFN-I production, the PR and VYR antiviral assays suggest the compounds do not act via IFN because these experiments were performed in Vero cells that do not produce active IFN-I.

Interestingly, while amino acids 222, 248, and 251 are conserved in all species of ebolavirus (5, 48), the residue 225 in Reston virus (RESTV) and Sudan virus (SUDV) is a lysine, and in Marburg virus (MARV) is an alanine. The retention of a basic amino acid in RESTV and SUDV demonstrates some conservation in VP35 interaction with Ub, albeit at reduced levels (see Table 1 and Figure 4C). However, the R225A mutation in MARV suggests reduced Ub binding and reduced polymerase activity. Overall, this points to the possibility that ebolaviruses are undergoing evolutionary changes on VP35 via interactions with the Ub system, although it is still unclear its relationship with pathogenesis since MARV is also highly pathogenic.

We have so far been unable to identify the E3-Ub ligase responsible for the production of these unanchored polyUb chains. It seems unlikely that TRIM6 is the E3 ligase involved in this process because we have shown TRIM6 can covalently modify VP35 (4), and can also synthesize unanchored K48-linked polyUb chains (35). In addition, the mechanisms by which covalent ubiquitination on K309 and binding of unanchored Ub seem distinct since K309 is located in the central basic patch and its ubiquitination promotes binding to L leading to regulation of viral transcription (6), while binding of unanchored Ub to the first basic patch of VP35 is most likely regulating binding with NP (11).

The function of unanchored Ub continues to be controversial. It has been proposed to regulate different immune pathways and inflammatory responses by the host (49). For example, we found that unanchored K48-linked polyUb regulates the antiviral response by promoting IKKε activation to stimulate IFN-I responses (35), and K63-linked polyUb chains can promote RIG-I activation (32, 34, 50–52), and TAK1 activation (36, 53, 54). On the other hand, viruses have been shown to utilize both unanchored Ub and covalent ubiquitination to replicate. In relation to unanchored Ub, Influenza viruses contain unanchored Ub in infectious virions, which can enhance virus replication by promoting uncoating via histone deacetylase 6 (HDAC6) (55, 56), and other viruses may use similar mechanisms (57). We have also detected Ub chains in sucrose-purified infectious EBOV, which could be covalently attached to VP35 or other viral proteins; however, Ub corresponding to low molecular weight chains suggest some unanchored Ub may be present (data not shown). At the moment, it is unclear if these Ub chains would act at the level of virus entry or would prime VP35 to initiate it is polymerase co-factor activity early during infection. This further supports a role for Ub in promoting early stages of virus replication.

Despite the existence of approved treatments for patients infected with EBOV, which include monoclonal antibodies mAb114 and REGN-EB3 (58, 59), outbreaks of EBOV in Africa continue to present a public health threat. Therefore, there is still a need for the identification of accessible treatments. In conclusion, we have used a computational approach combined with the identification of chemical compounds that block non-covalent VP35-Ub interactions to study functional outcomes. This approach exemplifies how computational docking can contribute to the study of host-virus interactions and predict potential therapeutic approaches.

## Materials and Methods

### Plasmids and Reagents

The VP35 constructs (VP35 R225E and R225K mutations) in the pCAGGS backbone were kindly provided by Dr. Basler (Icahn School of Medicine at Mount Sinai, NY), and were previously described (11). The mutations on VP35 K309R and K309G were previously described in (6). The IsoT plasmid was described in (35). The mutant plasmid sequences were confirmed using Sanger sequencing (UTMB Molecular Genomics). The other plasmids including Renilla luciferase, pCAGGS empty vector, and minigenome components (EBOV L, EBOV NP, EBOV VP30, T7 polymerase, and EBOV minigenome firefly luciferase) have been previously described (4, 6).

### Cells and viruses

Cells were obtained from the American Type Culture Collection (Manassas, VA). Vero cells (ATCC CCL81), and Vero E6 were used for infection studies, and HEK293T cells (ATCC CRL-3216) were used for transfection. Cells were maintained in 1X Dulbecco’s modified eagle’s medium (DMEM) (Gibco by Life Technologies, Inc., USA), supplemented with 5% Fetal Bovine Serum (FBS) low in endotoxins and heat-inactivated (Life Technologies, Inc., Rockville, MD, USA) and incubated at 37°C, with 5% CO2. Wild-type Ebola virus (EBOV) strain Zaire was propagated in Vero E6 cells. Virus titer was determined by plaque assay using Vero-CLL81 cells (6). All experiments performed with live EBOV were carried out in the Galveston National Laboratory Biological Safely Level 4 laboratory at The University of Texas Medical Branch at Galveston.

### Modelling the non-covalent interaction between VP35 and Ubiquitin

Protein-protein docking of Ub on VP35 was done using ClusPro 2.0 (60) and ZDOCK (61) web servers with default parameters, using Protein Data Bank (PDB) (62) entry IDs 1ubq and 3jke respectively. The top 10 poses from ZDOCK were kept for minimization. ClusPro divides the results into 4 categories: balanced, electrostatics-favored, hydrophobics-favored, and Van der Walls + Electrostatics favored. The top 5 poses from the balanced category and the top 3 from each other were kept (14 total from ClusPro).

These 24 poses were then submitted to minimization, equilibration and short 100 ps production molecular dynamics trajectories using GROMACS (63) version 2016.3, with parameters as described in the lysozyme tutorial from (64). The conformations and energies from the trajectories were saved every 2 ps. 3 replicate trajectories were computed for every of the 24 poses. The average total potential energies from these short runs were used to rank the different poses, and the top 4 poses were selected this way. These 4 poses were then simulated for 3 replicates of 1ns trajectories, using the same parameters as before.

The resulting trajectories were analyzed using the gRINN software (65) to find pairs of residues of low interaction energy at the protein-protein interface. We used default parameters with a stride of 10 (50 frames from each replicated 1 ns trajectory were analyzed). We were aiming to find pairs common to all docking poses and all frames, but surprisingly there were none. Instead, we chose to analyze only the 3 replicate trajectories from the lowest energy pose, according to the potential energy computed from the MD simulations (pose 4 from ZDOCK). This allowed us to rank all VP35-Ub interface pairwise residue interactions in decreasing order of relative contribution to the predicted binding free energy (ΔG).

We employed Surfaces (66), a software to quantify and visualize interactions within and between proteins and ligands. With Surfaces, we were able to evaluate the change in ι1G expected from each of the 19 mutations on VP35 for the top-ranked interacting position.

### Modelling Interactions between VP35, Ubiquitin and dsRNA

We modeled the complex involving VP35, Ubiquitin, and dsRNA using our modeled complex of VP35 and Ub as a template upon which we superimposed experimental structures of VP35 in complex with dsRNA (PDB entry 3KS8) and K63-linked Ubiquitin dimer (PDB entry 2JF5). In order to model a chain of three K63-linked Ub monomers, we used twice the 2JF5 structure, and superimposed the first chain of the dimer once and the second chain onto the Ub once. We utilized the Yasara energy minimization web server (67) to resolve any steric clashes in the ternary complex between the VP35-Ub complex and dsRNA. The minimized structure was further validated for the absence of steric clashes using the dedicated function provided by Surfaces scripts (66).

### Identifying potential protein-protein interaction disruptors of VP35-Ubiquitin through ultra-massive virtual screening

We used GetCleft, a C implementation of the Surfnet algorithm (68) to identify the top 5 largest cavities of VP35 using the structure generated by molecular dynamics. From these, we selected the cavities in contact with residues in the interface with Ub with the rationale that a molecule occupying such a cavity may disrupt binding to Ub. For the docking experiments, we selected molecules from the Chemical Component Dictionary (69) representing all ligands in complex with a protein present in the PDB. We selected all compounds composed of more than 4 non-hydrogen atoms of which 2 are carbons, for a total of 36,000 compounds. We utilized the ligand protein docking software FlexAID (70) to perform 10 docking simulations (500 generations and 500 chromosomes, or 250,000 pose evaluations) for each of the 36,000 molecules and ranked the molecules based on the mean docking score value (CF score). We selected the top 10% molecules for a second round of 10 docking simulations of 1,000 generations and 1,000 chromosomes (1×10^6^ evaluations). We proceeded to compare the level of binding-site similarities between the binding-site defined by each of the 20 top-ranked molecules and their known binding-site as seen in the PDB. For that, we utilized the IsoMIF (71) method for the detection of molecular interaction field similarities. The two compounds ultimately selected for experimental testing were molecules with high values of binding-site similarities and high docking scores but also favorable pharmacological properties.

### Plaque Reduction (PR) assay

24 hours before infection, confluent Vero CCL-81 cells were plated in a 12-well plate with 1X DMEM and 5% FBS and maintained at 37°C with 5% CO_2_. The EBOV inoculum was prepared in 2% FBS DMEM and 100μl inoculum/well was used with a final MOI of 0.01. Cells were incubated with the EBOV for 60 min at 37°C, 5% CO_2_ with constant gentle rocking. The EBOV inoculum was then removed, and cells were washed with PBS 1X and 2 mL of overlay (0.5% methylcellulose, diluted 1:1 with 2X MEM supplemented with 2% FBS, 1% penicillin/streptomycin and the corresponding compound concentration 200, 100, 50 and 25µM) were added each well. The EBOV and cell control was run parallel for each tested compound. Further, favipiravir (AdooQ^®^ Biosciences) was used as a positive control drug using the same experimental set-up as described for EBOV and cell control. Cells were incubated at 37°C with 5% CO_2_ for 12 days later the overlay was removed, and plates stained with 0.05% crystal violet in 10% buffered formalin for approximately twenty minutes at room temperature. The plates were then washed and dried, and the number of plaques were counted. The number of plaques in each set of compound dilutions were converted to a percentage relative to the untreated virus control. The 50% effective (EC_50_, virus-inhibitory) concentrations were then calculated by linear regression analysis. The cytotoxicity assay (In vitro Toxicology Assay Kit, Neutral red based; Sigma) was performed in 96-well plates following the manufacturer’s instructions. Briefly, the growth medium was removed from confluent cell monolayers and replaced with fresh medium (total of 100µl) containing the test compound with the concentrations indicated for the PR assay. Control wells contained medium with the positive control or medium devoid of the compounds. Wells without cells and growth medium only were used as blank. A total of up to five replicates were performed for each condition. Plates were then incubated for 12 days at 37°C with 5% CO_2_. The plates were then stained with 0.033% neutral red for two hours at 37°C in a 5% CO_2_ incubator. The neutral red medium was removed by complete aspiration, and the cells were rinsed 1X with phosphate-buffered solution (PBS) to remove the residual dye. The PBS was completely removed, and the incorporated neutral red was eluted with 1% acetic acid/50% ethanol for 30 minutes. The dye content in each well was quantified using a 96-well spectrophotometer at 540 nm wavelength and 690 nm wavelength (background reading). The 50% cytotoxic (CC_50_, cell-inhibitory) concentrations were then calculated by linear regression analysis. The quotient of CC_50_ divided by EC_50_ gives the selectivity index (SI_50_) value. The positive control compound was evaluated in parallel in each test.

### Virus yield reduction (VYR) assay

The VYR assay involves a similar methodology to the PR assay described above. Briefly, Vero CCL81 cells in a confluent monolayer were infected as described above. At the end of the incubation, the inoculum was removed and replaced with a fresh medium containing the compounds (200, 100, 50 and 25µM) and incubated at 37°C with a 5% CO_2_ for 72 hours. The supernatant was collected and used to determine the compound’s inhibitory effect on virus replication. The EBOV that was replicated in the presence of the compounds is titrated and compared to the EBOV levels in untreated, infected controls. For titration of supernatant, serial ten-fold dilutions were prepared and used to infect fresh monolayers of cells. Cells were overlaid with 0.5% methylcellulose, diluted 1:1 with 2X MEM as previously described, and the number of plaques was determined. Test compounds and positive controls were tested in triplicates. The viral titer in each set of compound dilutions was converted to a percentage relative to the untreated virus control. Plotting the log10 of the inhibitor concentration versus the percentage of virus produced at each concentration allows calculation of the 90% (one log10) effective concentration by linear regression. Cytotoxicity is determined in uninfected cells as described, and 50% cytotoxic (CC50) concentrations were then calculated by linear regression analysis. The positive control compound is evaluated in parallel in each test.

### Transfections and Immunoprecipitations

HEK293T cells were plated in 6-well plates (400,000 cells/well) in DMEM 1X supplemented with 5% FBS for 24 hours, followed transfection with the specific plasmid (Flag-VP35 WT, Flag-VP35 R225E, Flag-VP35 R225K, Flag-VP35 K309R, Flag-VP35 K309G, Flag-VP35 N-terminus, Flag-VP35 C-terminus, HA-Ub WT, HA-Ub ΔGG, His-IsoT WT, His-IsoT C335A and pCAGGS empty vector), using TransIT-LT1 (Mirus) following the manufacturer’s recommendations. 30 hours after transfection, cells were lysed in RIPA buffer containing complete protease inhibitor (Roche), n-ethylmaleimide (NEM), and iodoacetamide (IA) (RIPA complete). Lysates were cleared at 25,200 xg for 20 minutes at 4°C, and 10% of the clarified lysate was added to 2X Laemmli sample buffer (BioRad) with 5% ß-mercaptoethanol and boiled at 95°C for 10 minutes to generate whole cell extracts (WCE). The remaining clarified lysate was mixed with 10uL of anti-Flag-Agarose or anti-HA-Agarose beads (Sigma) and incubated at 4°C overnight on a rotating platform. Later, the beads were washed seven times with RIPA buffer (SDS0.1% (v/v), Deoxycholic acid sodium salt 0.5% (w/v), Tris Ph 8.0 50mM, NaCl 150mM, NP-40 1%) with IA and NEM, after the seven RIPA washes. If elution was not necessary, the beads were mixed with 50µl 2x buffer Laemli and boiled to 95°C for 10 minutes. For elution, seven RIPA washes, an additional wash made using peptide elution buffer (10mM Tris Ph 7.4 and 150mM NaCl in nuclease-free water (NF H2O)) without peptide were done. The protein was then eluted in 15 μL of peptide elution buffer three times using Flag peptide or HA peptide (Sigma) 300µg/mL. Finally, 4x buffer Laemli was added to each elution, and boiled to 95°C for 10 minutes.

To test Ubiquitin interactions, an additional wash using NT2 buffer (Tris 7.4 50mM, NaCl 150mM, MgCl 1 mM, NP-40 0.05% (v/v)) was done, and the beads were kept with 200 µl of NT2 buffer and added K63 polyUb chains (2-7 chains) (UBPBio) and incubated at 4°C overnight on a rotating platform. 10% of these was added to 2X Laemmli sample buffer (BioRad) with 5% ß-mercaptoethanol and boiled at 95°C for 10 minutes to generate the inputs. The beads were washed seven times with NT2 buffer and elution was made as described previously.

To test Ubiquitin interactions with VP35 and small compounds, immunoprecipitations were done following a similar protocol to test Ubiquitin interactions. Briefly, an additional wash using NT2 buffer was performed and the beads were incubated with 200 µl of NT2 buffer and treated with 3-[4-(aminosulfonyl) phenyl]propanoic acid (pCEBS) (Enamine US inc.) or 2,5-Dimethyl-4-sulfamoyl furan-3-carboxylic acid (SFC) (Enamine US inc.) pCEBS and SFC were diluted to final concentrations of 200, 100, 50, 25 and 12µM in 1% dimethyl sulfoxide (DMSO). The beads were incubated at room temperature (RT) for 1 hour on a rotating rocker and were incubated with K63 polyUb chains (2-7 chains) (UBPBio) at 4°C overnight on a rotating platform. 10% of these were added to 2X Laemmli sample buffer (BioRad) with 5% ß-mercaptoethanol and boiled at 95°C for 10 minutes to generate the inputs. Beads were washed seven times with NT2 buffer, and elution was done as mentioned previously.

### Minigenome assay

The monocistronic minigenome construct expressing the firefly luciferase gene (72) was kindly provided by Dr. Bukreyev (UTMB). The plasmids pCEZ-NP, pCEZ-VP35, pCEZ-VP30, pCEZ-L, and pC-T7 were previously described (73). 293T cells were plated (50,000 cells/well) onto 24-well plates in 5% FBS 1X DMEM for 24 hours, and co-transfected with the following plasmids: EBOV minigenome (125 ng), pCEZ-VP30 (31.25 ng), pCEZ-NP (62.5 ng), pCEZ-L (500 ng), pC-T7 polymerase (125 ng), 100 ng of empty vector (pCAGGS) or pCAGGS-VP35 (WT, R225E, or R225K), and REN-Luc/pRL-TK plasmid (20 ng; Promega) expressing Renilla luciferase used as an internal control to normalize transfection efficiency. To test the compounds; four hours post-transfection, treatments with PCEBS or SFC to 200, 100, 50, 25, and 12µM in 1% DMSO were added, and cells were incubated at 37°C, 5% CO2 for fifty hours. After, the cells were lysed to measure the luciferase signal using the Dual-Luciferase Reporter Assay System (Promega) with a Cytation 5 reader (Biotek). A portion of the lysate was boiled at 95°C for 10 minutes in 2X Laemmli buffer to evaluate protein expression via immunoblot.

### Western blot

Protein samples were run on 4–15% or 7.5% Criterion-TGX Precast Gels (Bio-Rad). The proteins were transferred onto a methanol-activated Immun-Blot PVDF membrane (Bio-Rad), and the membrane was blocked in 5% Carnation powdered skim milk (Nestle) in 1X TBS-T (20mM Tris, 150mM NaCl) for 1 hour. Primary antibodies were prepared in 3% bovine serum albumin 1X TBS-T with 0.02% sodium azide to the appropriate dilution: anti-Flag and anti-HA (Sigma) 1:2000, anti-VP35 (6C5 Kerafast) 1:1000, anti-NP (provided by Dr. Basler, Mount Sinai), anti-ubiquitin (Enzo) 1:1000, and anti-β-actin (Abcam) 1:2000. The next day, the blot was washed three times every 5 min using 1X TBS-T before incubation with HRP-conjugated goat-anti-rabbit (GE Healthcare) or goat-anti-mouse (GE health care) for 1 hour. The blot was washed in 1X TBS-T and developed using Pierce ECL Western Blotting Substrate (Thermo Fisher) or SuperSignal West Femto Maximum Sensitivity Substrate (Thermo Scientific). For blot quantifications, the area under the curve (AUC) was measured for each band of interest using ImageJ.

### Statistical analysis

Three independent assays were carried out in triplicate for a single-factor analysis of variance to determine if differences exist among the groups: treated, positive controls, negative controls, and infection. A Shapiro–Wilk test was performed to determine normality in the data. Multiple comparison tests were performed to determine among which groups statistically significant differences are occurring compared to the infection control using Tukey’s multiple comparison test. These tests were run through the GraphPad Prism v.9 statistical package (San Diego, CA, USA). All tests were considered statistically significant when *p* < 0.05 with 95%CI.

## Supporting information

Supplementary Pymol Session File 1

## Acknowledgments

This work was supported by NIH/NIAID grants R01AI134907 to M.I.G and R.R, R01AI166668, R01AI155466, P01AI150585 awarded to R.R., T32AI060549 awarded to S.v.T. RJN is a member of the Quebec network for the study of protein structure, function and engineering (PROTEO).

## SUPPLEMENTARY DATA

Supplementary table 1 shows the predicted contributions of individual interactions in kcal/mol to the binding energy between VP35 and Ubiquitin calculated by gRINN and Surfaces. The first row in blue shows the residue ARG225, predicted as the highest and second highest contributing favorable interaction respectively by gRINN and Surfaces.

Supplementary table 2 shows the difference in predicted contributions of individual interactions to the ΔG of binding for different mutants in position 225. In blue in the first row the original ARG residue. Note that the value for the interaction with GLUE18 alone is the same as what is presented in Supplementary Table 1. The second value (overall), represents the sum of all interactions with the specific residue. For example, R225 makes additional favorable interactions with other residues that contribute to the total energy of binding. The values in salmon highlight the predictions for the mutations for K225 and E225.

**Supplementary Table 1.** Comparison of contribution to binding energy of gRINN and Surfaces predictions in kcal/mol of individual interactions within the VP35-Ub complex interface.

| VP35 | Ub | gRINN | Surfaces |
| --- | --- | --- | --- |
| ARG225 | GLU18 | -14.81 | -1.11 |
| LYS222 | GLU16 | -14.00 | 0.00 |
| ARG305 | ASP58 | -10.60 | -0.75 |
| ARG298 | GLU24 | -10.43 | -1.85 |
| ASP230 | LYS63 | -9.00 | 0.00 |
| TYR229 | GLU18 | -4.45 | -0.59 |
| GLN244 | GLU18 | -1.99 | -0.01 |
| PRO304 | ASP58 | -1.65 | 0.00 |
| ARG305 | SER57 | -1.63 | -0.99 |
| LYS222 | GLU18 | -1.60 | 0.00 |

**Supplementary Table 2.** Predicted contributions of different residues modeled in position 225 to the binding energy in kcal/mol overall or specifically with GLU18.

| Mutation | Contribution to Binding Energy |  |
| --- | --- | --- |
|  | Overall | GLU18 |
| ARG225 | -1.27 | -1.11 |
| TRP225 | -0.81 | -0.81 |
| LYS225 | -0.57 | -0.47 |
| GLN225 | -0.56 | -0.56 |
| THR225 | -0.03 | -0.03 |
| ASN225 | 0.10 | 0.10 |
| LEU225 | -0.03 | -0.03 |
| VAL225 | -0.05 | -0.05 |
| ILE225 | -0.10 | -0.10 |
| MET225 | -0.12 | -0.12 |
| CYS225 | 0.00 | 0.00 |
| SER225 | 0.00 | 0.00 |
| PRO225 | 0.00 | 0.00 |
| GLY225 | 0.00 | 0.00 |
| HIS225 | 0.00 | 0.00 |
| ALA225 | 0.00 | 0.00 |
| TYR225 | 0.00 | 0.00 |
| GLU225 | 0.07 | 0.07 |
| PHE225 | 0.00 | 0.00 |
| ASP225 | 0.13 | 0.13 |

